# Emergence of microglia structural and functional heterogeneity between hippocampal subregions during development into early adulthood

**DOI:** 10.64898/2026.02.04.703826

**Authors:** Eric W. Salter, Emily Lackie, John Georgiou, Graham L. Collingridge

## Abstract

Phagocytosis is a key function performed by microglia to maintain tissue homeostasis. The degree of microglial phagocytic activity differs across ages and between gross anatomical brain regions, dictated by the local environment. Here, we asked whether microglial phagocytic phenotype exhibits subregion-scale tuning by circuit-specific features within a brain region. To address this, we took advantage of the highly stereotyped architecture of the hippocampus. We examined three adjacent synaptic subregions, the CA1 stratum radiatum (SR), stratum lacunosum moleculare (SLM) and dentate gyrus molecular layer (DGML). These three subregions provide an ideal system for examining local microglia heterogeneity, as each subregion contains distinct neuropil features, creating three adjacent unique micro-environments to which the microglia are exposed. We measured the phagocytic activity and morphological properties of over 1,000 individual microglia at two developmental points, mid-postnatal (P16) and early adulthood (P60) in the CA1 SR, SLM and DGML. We found that microglial phenotype diversified with development into early adulthood. At the mid-postnatal age, phagocytic activity and morphology were homogeneous across subregions. Conversely, in young adulthood, microglia in the CA1 SR and DGML exhibited a reduction in phagocytic activity, while microglia in the CA1 SLM maintained a highly phagocytic phenotype reminiscent of an immature-like state. These findings uncover a fine-scale tuning of microglia activity that emerges during maturation and is dictated at the sub-region level of the hippocampus, uncovering a distinct population of microglia in the CA1 SLM that exhibit a persistent immature phenotype under physiological conditions. Understanding the target(s) of this phagocytosis and consequences for CA1 SLM function will provide new insight into the role of local tuning of microglia properties for circuit-specific needs in both health and disease.

## INTRODUCTION

Microglia, the principal immune cells in the brain parenchyma, exhibit substantial phenotypic diversity in both space and time due to distinct local environmental signals (Matcovitch-Natan et al., 2016; De Biase et al., 2017; Ayata et al., 2018; De et al., 2018; Böttcher et al., 2019; Hammond et al., 2019; Li et al., 2019; Masuda et al., 2019; Stratoulias, Venero, Tremblay, & Joseph, 2019; Masuda et al., 2020; Tan, Yuan, & Tian, 2020; Young et al., 2021; Barko et al., 2022; Colombo et al., 2022; De Felice et al., 2022; Paolicelli et al., 2022; Stratoulias et al., 2023; van Weering, Nijboer, Brummer, Boddeke, & Eggen, 2023). For example, cell density and morphology are dependent on when and where microglia are located in the brain (Lawson et al., 1990; Savchenko, McKanna, Nikonenko, & Skibo, 2000; Grabert et al., 2016; Colombo et al., 2022; van Weering et al., 2023). The advent of single-cell omics technologies has been pivotal for further defining the microglia states and heterogeneity under physiological and pathological contexts (Matcovitch-Natan et al., 2016; Böttcher et al., 2019; Hammond, Dufort, et al., 2019; Li et al., 2019; Masuda et al., 2019; Masuda, Sankowski, Staszewski, & Prinz, 2020; Young et al., 2021). However, the full breadth of microglia diversity at the functional level remains to be uncovered.

There is evidence that microglia phenotype is more diverse in early development compared to adulthood due to the varied unique brain states specific to development (Wlodarczyk et al., 2017; Hammond, Dufort, et al., 2019; Li et al., 2019; Prinz et al., 2019; Shen, Qiu, Wight, Kim, & Cantor, 2022). Further, microglia are canonically considered to be most phagocytic at earlier ages because of the higher levels of neuron and synapse turnover in development (Schafer et al., 2012; Cunningham et al., 2013; Weinhard et al., 2018; Anderson et al., 2019; VanRyzin et al., 2019). Studies examining the spatial heterogeneity of microglia state and function have commonly used a broad segregation of brain regions (e.g. hippocampus, cortex, cerebellum, etc.) (Lawson et al., 1990; Greter et al., 2012; Grabert et al., 2016; Verdonk et al., 2016; Böttcher et al., 2019; Li et al., 2019; Masuda et al., 2019; Stratoulias et al., 2019; Tan et al., 2020; Young et al., 2021). However, whether heterogeneity of microglia phagocytic function exists between local circuits within the same brain region under homeostatic conditions, and how such heterogeneity evolves over development, is not known.

The hippocampus is a well-suited brain region to address this gap in knowledge, as it is composed of subregions with discrete borders delineating distinct synaptic and cellular compositions (Klausberger & Somogyi, 2008; Cembrowski & Spruston, 2019; Topolnik & Tamboli, 2022). This is exemplified by pyramidal neurons in the CA1: excitatory inputs from upstream hippocampal CA3 synapse onto proximal dendrites in the stratum radiatum (SR), with a sharply defined border for long-range excitatory inputs primarily from the entorhinal cortex (EC) onto distal dendrites in the stratum lacunosum moleculare (SLM; van Groen et al., 2003; Klausberger & Somogyi, 2008; van Strien et al., 2009). Thus, microglia in the CA1 SR exist in a distinct synaptic environment compared to microglia in the CA1 SLM.

Here, we took advantage of this anatomical segregation to define the spatiotemporal heterogeneity of microglia in the hippocampus across subregions and development. We discovered that while microglial phagocytosis and morphology are homogenous between subregions in postnatal mice, an immature-like phagocytic and morphological state specifically in the CA1 SLM persists into adulthood. Thus, unlike global principles of microglia heterogeneity, at the subregion level in the hippocampus, microglial functional heterogeneity increases during development into adulthood. These findings establish the hippocampus as an ideal model system to study the mechanisms and implications of local tuning of microglia function in health and disease.

## RESULTS

To assess hippocampal subregion heterogeneity of microglia phagocytic activity, we examined the CA1 SR and CA1 SLM due to the distinct pre-synaptic inputs onto the same postsynaptic CA1 pyramidal neurons. We assessed a third subregion, the dentate gyrus molecular layer (DGML), as the EC also sends long-range excitatory inputs to the DGML in addition to the CA1 SLM, synapsing onto post-synaptic dentate gyrus granule cells. Thus, by analyzing these three hippocampal subregions (CA1 SR, SLM and DGML) we can assess microglia phenotype in local environments with distinct combinations of pre- and post-synaptic partners (Figure 1A).

**Figure 1.**
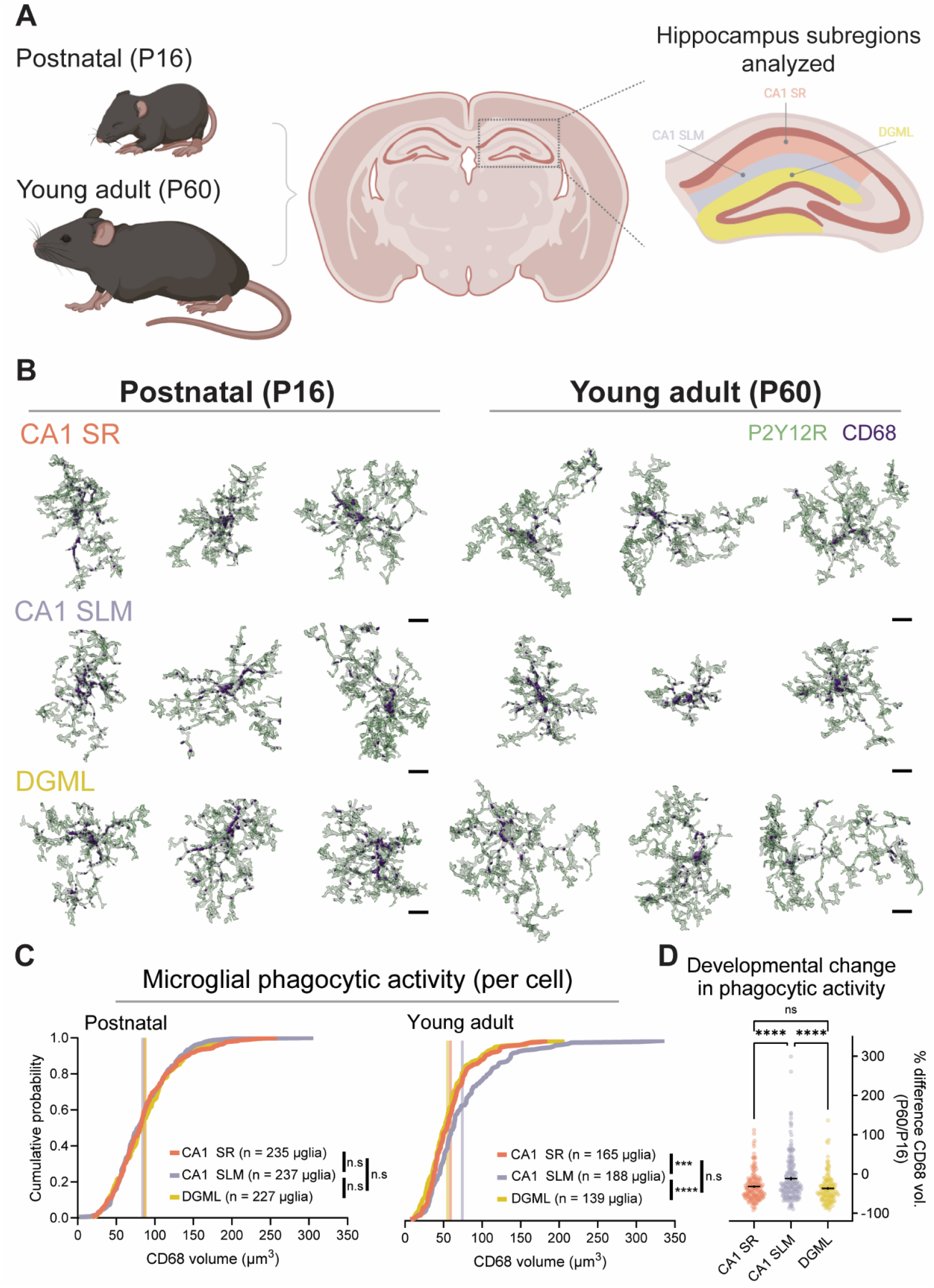
CA1 SLM microglia retain high phagocytic activity into early adulthood. **A** Schematic of age groups and hippocampal subregions analyzed in the current study. **B** Representative single-cell renderings of the lysosome marker CD68 (solid purple) within P2Y12R (transparent green) to delineate microglial membrane, across the CA1 SR (*upper*), CA1 SLM (*middle*) and DGML (*lower*) in postnatal (*left*) and young adult (*right*) mice. Scale bars are 8 µm. **C** Cumulative probability distributions for postnatal (*left*) and young adult (*right*) mice of cumulative CD68 volume within individual microglia (microglial phagocytic activity) across the CA1 SR, CA1 SLM and DGML. Vertical lines denote the mean value for each subregion. In the postnatal age group, there was no significant difference in CD68 volume per microglia between hippocampal subregions. In the young adult age group, total CD68 volume in CA1 SLM microglia was significantly higher compared to the CA1 SR and DGML. **D** Quantification of percent difference in the total CD68 volume per microglia in young adults compared to the postnatal average in each hippocampal subregion. Microglial CD68 volume was reduced to a significantly higher percentage in the CA1 SR and DGML compared to the CA1 SLM.

We assessed two ages, late postnatal (P16) and early adulthood (P60) in both male and female mice (Figure 1A). These two time points were chosen as microglial numbers peak in the hippocampus at this postnatal timepoint (Kim et al., 2015; Weinhard et al., 2018), indicating this is an important time for microglia function in the developing hippocampus. The adult age was chosen as previously it has been found that microglia adopt a stable, mature transcriptional state by this time (Matcovitch-Natan et al., 2016). Microglia were delineated by immunofluorescence (IF) staining using the microglia-specific transmembrane protein P2Y12R (Sasaki et al., 2003; Haynes et al., 2006; Hickman et al., 2013; Y. Zhang et al., 2014; Bosco et al., 2018; McKinsey et al., 2020). We co-stained brain tissue for the myeloid lysosome marker CD68, which is widely used to assay the extent of ongoing phagocytic activity of microglia (Schafer et al., 2012; Schafer, Lehrman, Heller, & Stevens, 2014; De Biase et al., 2017; Ayata et al., 2018; Kopec, Smith, Ayre, Sweat, & Bilbo, 2018; Scott-Hewitt et al., 2020). Volumetric confocal imaging followed by 3D reconstruction of individual microglia (see Methods) enabled us to analyze the spatiotemporal heterogeneity of microglia phagocytic activity at the single-cell level for nearly 1,200 microglia. We quantified microglia phagocytic activity as the total amount of CD68 within each cell (Figure 1B). In the mid-postnatal age group, there was no significant difference in microglia CD68 levels between the three subregions analyzed (Figure 1C), indicating that microglia phagocytic activity is homogenous across the CA1 SR, CA1 SLM and DGML at this age. Conversely, in the young adult age group, microglia in the CA1 SLM contained significantly more CD68 compared to both the CA1 SR and DGML (Figure 1C). Further, microglial CD68 levels were not significantly different between the CA1 SR and DGML in young adults (Figure 1C), indicating a distinct phagocytic phenotype of CA1 SLM microglia.

We next asked whether the subregion heterogeneity in phagocytic activity that emerged in young adults was the result of enhanced phagocytic activity compared to development in the CA1 SLM or maintenance of postnatal levels of phagocytic activity selectively in this subregion. To answer this, we normalized the CD68 levels of individual microglia in young adults to the postnatal mean in each hippocampal subregion. This analysis revealed that microglial phagocytic activity was reduced to a significantly greater degree in the CA1 SR and DGML compared to the CA1 SLM of young adults (Figure 1D). These results together elucidate that microglia specifically in the CA1 SLM retain an immature-like level of phagocytic activity under homeostatic conditions. Microglia structure is tightly linked to function (Prinz et al., 2019; Stratoulias et al., 2019; Paolicelli et al., 2022). Therefore, we assessed whether the morphology of microglia also exhibited emergent heterogeneity across hippocampal subregions. We performed 3D filament reconstruction of individual cells identified through P2Y12R staining, and analyzed two key morphological metrics: i) total branch length reflecting microglia territory, and ii) number of branch points, which measures the morphological complexity of cells (Figure 2A). Similar to phagocytic activity, in postnatal developing mice neither total microglia branch length nor number of branch points was significantly different between any of the three hippocampal subregions (Figure 2B,C). However, in young adult mice, both total microglia branch length and number of branch points were significantly lower in the CA1 SLM compared to both the CA1 SR and DGML. Further, DGML microglia from young adult mice exhibited significantly higher branch length and number of branch points compared to CA1 SR microglia (Figure 2B,C). These results indicate that the immature-like phagocytic state of young adult CA1 SLM microglia is accompanied by reduced morphological territory and complexity. The results of our morphology analysis provide additional support for the emergence of microglia phenotype heterogeneity between subregions in adulthood.

**Figure 2.**
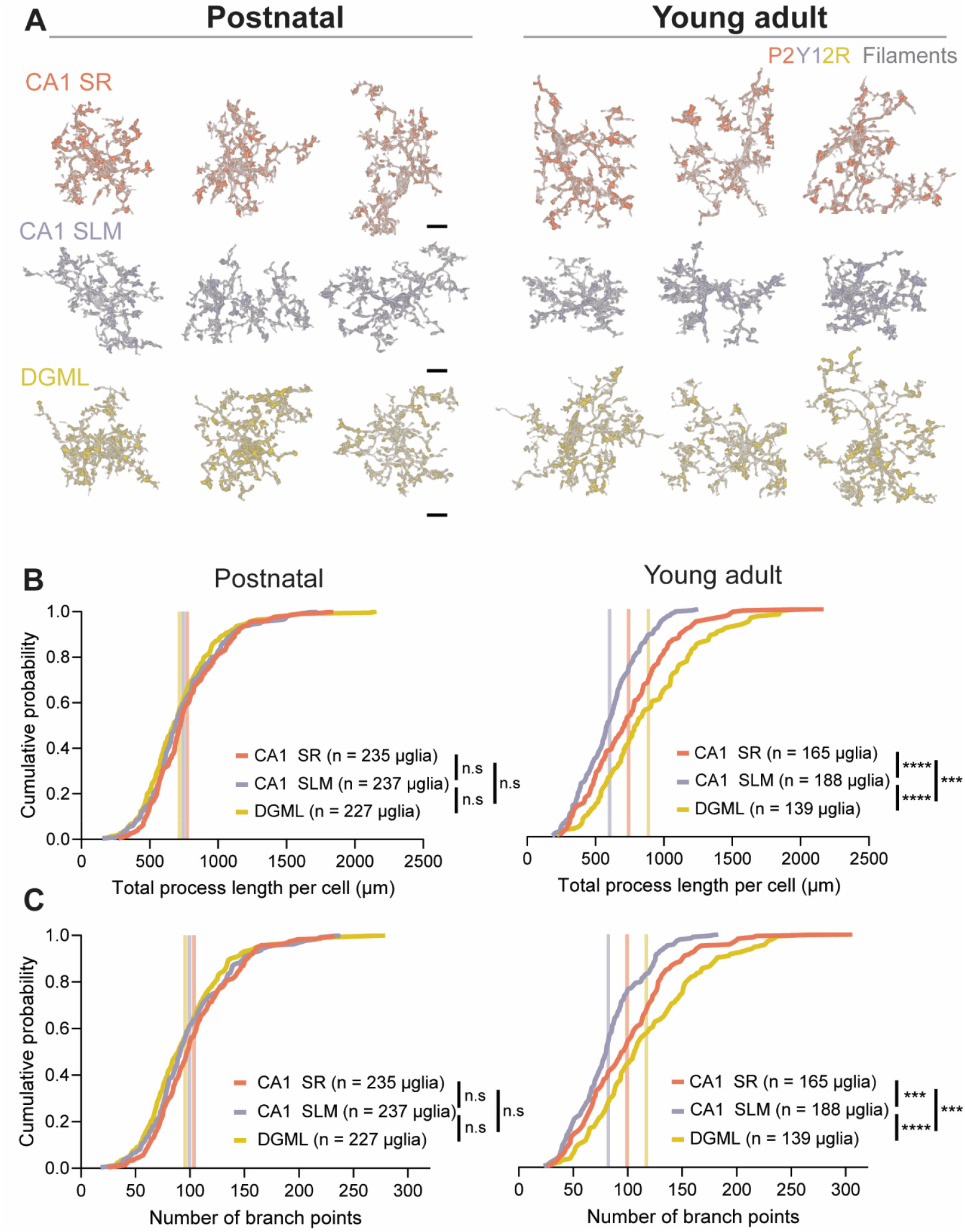
Heterogeneity of microglia morphology between hippocampal subregions emerges in early adulthood. **A** Representative filament tracer overlays in P2Y12R intensity images colour-coded based on hippocampal subregion in postnatal (*left*) and young adult (*right*) mice. Scale bars are 8 µm. **B** Cumulative probability distribution for total microglia branch length of individual microglia in the CA1 SR, CA1 SLM and DGML. Summed microglia branch length was not significantly different between hippocampal subregions in the mid-postnatal age group. In the young adult age group, microglia branch length was significantly lower in the CA1 SLM compared to the CA1 SR and DGML. Branch length was also significantly higher in the DGML compared to the CA1 SR. Vertical lines denote subregion mean value. **C** As per **B** but for number of filament branch points. The number of branch points of CA1 SLM microglia was significantly lower than CA1 SR and DGML microglia at P60. Additionally, DGML microglia had significantly more branch points compared to CA1 SR microglia at the young adult stage.

## DISCUSSION

Phagocytosis of cellular debris is a primary homeostatic function of microglia under physiological conditions (Galloway et al., 2019; Márquez-Ropero, Benito, Plaza-Zabala, & Sierra, 2020; Borst, Dumas, & Prinz, 2021). There is an increasing understanding that the degree and targets of microglial phagocytosis are diverse across brain regions. Studies examining microglia heterogeneity have used gross anatomical segregation of brain regions and have therefore examined the hippocampus as a singular structure (Lawson et al., 1990; Greter et al., 2012; Grabert et al., 2016; Verdonk et al., 2016; Böttcher et al., 2019; Li et al., 2019; Masuda et al., 2019; Tan et al., 2020; Young et al., 2021). This broad delineation overlooks the diverse local circuits that exist within the hippocampus which may dictate distinct microglia states. Here, we utilized the stereotyped laminar architecture of the hippocampus to determine whether microglia phenotype is dictated by subregion cues. In postnatal development, microglia phagocytic activity and morphology were homogenous across the CA1 SR, CA1 SLM and DGML. However, in early adulthood, we uncovered that microglia specifically in the CA1 SLM maintain an immature phenotype. Previous studies examining brain-wide microglia heterogeneity found that microglia states become less heterogeneous during development (Hammond et al., 2019; Li et al., 2019). Our findings reveal that locally in the hippocampus, microglia heterogeneity instead emerges as mice mature. Thus, the rules that govern microglia state transitions differ at the global vs. local level.

Two important questions that arise from our findings are: i) what are the CA1 SLM-specific cues driving the distinct microglia state, and ii) which microglia receptor(s) are responsible for receiving these cues. While the post-synaptic excitatory neurons in the CA1 SR and SLM are the same (pyramidal neurons), it is possible that phagocytic-inducing cues are trafficked selectively to distal dendrites in the SLM. Alternatively, pre-synaptic inputs could be responsible for inducing the SLM-specific microglia state. The most extensively studied input to this subregion are the long-range excitatory inputs from the EC (van Groen et al., 2003; Klausberger & Somogyi, 2008; van Strien et al., 2009). Although the EC also innervates the DGML (where we found that microglia do not exhibit the same immature state) the CA1 SLM is preferentially targeted by layer III EC neurons, while the DGML is targeted primarily by layer II EC neurons (van Strien et al., 2009; Suh et al., 2011). As such, highly specific pre-synaptic cues on EC inputs only to the CA1 SLM could be responsible. The CA1 SLM also contains unique inhibitory circuits compared to the CA1 SR and DGML. Neurogliaform cells are a subtype of interneuron with somas and axonal arbors largely restricted to the CA1 SLM (Capogna, 2011; Bezaire & Soltesz, 2013; Overstreet-Wadiche & McBain, 2015). Additionally, somatostatin^+^ oriens-lacunosum moleculare interneurons selectively innervate the CA1 SLM (Freund & Buzsáki, 1996; Maccaferri & Lacaille, 2003; Maccaferri, 2005). Interestingly, somatostatin^+^ interneuron boutons are phagocytosed by microglia in the cortex (Gesuita et al., 2022), making these interneurons an attractive candidate for being the target of CA1 SLM microglia phagocytosis. A final possibility is that the cues are derived from non-neuronal cells. For example, microglia have been found to engulf myelin sheaths (Hughes & Appel, 2020) and myelination is highly enriched in the CA1 SLM compared to both the CA1 SR and DGML in mice (Park et al., 2016; DeFlitch et al., 2022). An ever-expanding list of microglial receptors linked to phagocytosis (Beiter, Sheehan & Schafer, 2024) suggests that the same endpoint can be achieved by multiple mechanisms, likely depending on the target. To this end, spatial-omics (Liu et al., 2024) may enable identification of candidate receptors/pathways enriched in CA1 SLM microglia compared to CA1 SR and DGML microglia to provide a mechanistic understanding of our observed phenotype. Finally, in the CA1 SLM of young adult mice, microglia are phenotypically immature. Future studies are required to dissect whether this is due to continuous presence of the same drivers/cues only in the CA1 SLM, or whether unique cues that generate same phenotype as development arise in early adulthood in this subregion.

Together, our findings elucidate that microglia in the hippocampus cannot be considered as a single entity under physiological conditions, and instead their functional phenotype is driven by the local circuit in which they are embedded. The findings may be relevant to disorders with regional hippocampal vulnerability, notably multiple sclerosis, psychiatric disorder, tauopathies and Alzheimer’s disease (Sicotte et al., 2008; Maurin et al., 2014; Michailidou et al., 2015; Nakahara et al., 2018; Planche et al., 2018), in which myelin and synaptic elimination by microglia occurs (Beiter, Sheehan & Schafer, 2024). Our study also establishes the CA1 SLM as a new model system to examine the determinants and functions of fine-scale microglia heterogeneity.

## METHODS

### Mice

All experiments and procedures were approved by The Centre for Phenogenomics (TCP) animal care committee and conformed to the Canadian Council on Animal Care (CCAC) guidelines. Mice were allowed *ad libitum* access to food and water, and were maintained on a 12:12 hour light:dark schedule (lights on at 7:00 AM). Mice were maintained on a C57BL/6J background. Equal numbers of male and female mice were used for each group.

### Brain fixation

Tissue was harvested using transcardial perfusion followed by post-immersion fixation. Mice were deeply anesthetized with isoflurane and were perfused at 6 mL/min, first with 15 mL of phosphate-buffered saline (PBS; 4^°^C) containing heparin (20 U/mL; Sigma, H3393) followed by 15 mL 4% paraformaldehyde (PFA) PBS (4^°^C) prepared fresh from 16 % stock (EM Sciences, 15710). Brains were then extracted, the frontal lobe and cerebellum were removed and then were post-fixed in 10 mL of 4% PFA PBS for 4.0 h at 4^°^C. Brains were then washed three times in PBS (15 min per wash) and then dehydrated in 20 mL of 30 % sucrose PBS (w/v), and kept at 4^°^C for 72 h. Finally, brains were embedded in tissue freezing medium (Tissue Tek, 4583 or EM Sciences, 72592), flash frozen on liquid nitrogen and stored at -80^°^C until sectioning.

### Immunofluorescence (IF)

Brains were sectioned at a thickness of 40 µm on a Cryostat (Leica) at -20^°^C and stored as free-floating sections at 4^°^C in PBS containing sodium azide (0.03%; Fisher Scientific, 71448-16) for up to 5 days prior to immunostaining. First, sections were washed 3 × 10 min to remove residual tissue freezing medium. Next, sections were incubated for 2 h in blocking/permeabilization buffer (PBS containing 0.2% Triton-X100 [MilliporeSigma, 9036-19-5], 0.03% sodium azide, 10% normal donkey serum [NDS; Jackson Immunoresearch, 017-000-001] and 50 mM glycine [MilliporeSigma, G7126]) at 4^°^C on a shaker. Subsequently, sections were transferred to primary antibody buffer (PBS containing 0.2% Triton-X100, 0.03% sodium azide, 10% NDS) overnight at 4^°^C on a shaker. The following primary antibodies and dilutions were used: Rat anti-CD68 (1:250, BioRad, MCA1957); Rabbit anti-P2Y12R (1:250, Anaspec AS 55043a). Sections were then washed 3 × 10 min followed by 3 × 30 min in PBS and then incubated in secondary antibody buffer (PBS containing 0.2% Triton-X100, 0.03% sodium azide, 10% NDS) overnight at 4^°^C on a shaker in the dark. The following secondary antibodies were used (all at a dilution of 1:100): Donkey anti-rabbit-Alexa488 (Jackson Immunoresearch, 711-545-152); Donkey anti-rat-Cy5 (Jackson Immunoresearch, 712-175-153). Sections were again washed 3 × 10 min followed by 3 × 30 min in PBS in the dark on a shaker. Finally, sections were adhered onto ethanol-cleaned coverslips (Thorlabs, CGKH15), dried, and then 100 µL of Prolong Glass (Invitrogen, P36980) was added to the sections, which were then mounted onto Superfrost plus microscope slides (Fisher Scientific, 22-037-246). Mounted sections were left to cure in the dark at RT for 48-60 h to improve index refraction matching between the mounting media and coverslip/objective oil.

### Confocal imaging

All confocal images were collected at the OptIma facility (NBCC - Network Biology Collaborative Centre Advanced Imaging Facility (RRID: SCR_025389) at the Lunenfeld-Tanenbaum Research Institute) using a Nikon A1R HD25 microscope and 60×, 1.4 numerical aperture (NA) oil immersion objective lens (Plan Apo). A 2.5 x zoom factor was used at 1024 x 1024 pixels (yielding a 110 nm pixel size) with 250 nm axial z-steps (9 µm total z-range). 3-4 technical replicate sections were used per mouse. Within each section, 2 fields of view per hippocampal subregion were imaged for each mouse. For each batch of technical section replicates, all mice from the analyzed cohort were imaged within a single day. All microscope settings were kept identical across technical replicate batches.

### Image analysis

All images were subject to identical image analysis steps. First, images were median filtered (3-pixel width for P2Y12R channel, 2-pixel with for CD68 channel) followed by rolling ball background subtraction (10 µm width for P2Y12R channel, 5 µm width for CD68 channel) using Nikon NIS elements. Single cell analysis of pre-processed images was then performed using Imaris (Bitplane). The P2Y12R channel was used to segment individual microglia. Imaris machine learning segmentation was trained on a subset of images, consisting of 2 images per subregion/age across two technical replicate batches. The machine learning segmentation was refined by experimenter input focusing on capturing fine microglia processes without over-segmenting somatic regions, which is a common issue when using intensity-based thresholding. This segmentation method also accounts better for minor technical replicate batch-to-batch intensity variations. The final trained model was then used to generate a P2Y12R surface in each image. To separate individual cells, surfaces were split using a seed point method. We empirically determined that using the CD68 channel to generate seed points (based on fixed intensity sum of squares) yielded the most consistent seeding of microglia somata compared to using the P2Y12R channel itself. This is because microglial lysosomes (marked by CD68) are largest and most dense at the soma, while the P2Y12R signal is diffuse across the entire membrane. Individual microglial P2Y12R surfaces were manually inspected and those that were artificially split during segmentation were rejoined manually. A CD68 surface was then generated using a fixed value intensity threshold. To analyze only CD68 within microglia, CD68 surfaces were filtered to only include those with an overlap ratio with the P2Y12R surface >0.3 and larger than 10 voxels to remove background. This microglial CD68 surface was used to generate final total CD68 surface volume per microglia. To quantify morphological parameters, Imaris filaments were generated for each microglia. First, the single cell P2Y12R surfaces were used to mask the P2Y12R intensity channel. Next, filaments were generated separately for each individual microglia using the masked intensity channel. Identical filament creation parameters were used for all cells. The seed point for the microglia soma was manually selected. The sum of the lengths of all filaments (total branch size) and the number of branching points were exported for analysis.

### Statistical analyses

All statistical tests were performed using GraphPad Prism (v10). Unless stated otherwise, a one-way ANOVA followed by *post hoc* Tukey’s multiple comparisons. * p <0.05, ** p <0.01, *** p<0.001, **** p < 0.0001.

## ACKNOWLEDGEMENTS

This work was supported by a CIHR (Canadian Institutes of Health Research) Foundation Grant to G.L.C. (#154276). G.L.C. is the holder of the Krembil Family Chair in Alzheimer’s Research. We thank the trainee support provided by the C.R. Younger Foundation. The work was supported by the MS Society of Canada as well as gifts from Jon and Nancy Love and the Dani Reiss Family Foundation (Neurodegeneration and Aging Research Program). E.W.S. was supported by a CIHR Doctoral Award (Frederick Banting and Charles Best Canada Graduate Scholarship, CGS-D) and an Ontario Graduate Scholarship (OGS). The NBCC imaging facility at LTRI is supported by the Canada Foundation for Innovation and the Ontario Government.

## AUTHOR CONTRIBUTIONS

E.W.S.: conceptualization, investigation, analysis, visualization, and writing; E.L.: image analysis. J.G.: supervision, editing, funding acquisition, project oversight. G.L.C.: conceptualization, supervision, editing, funding acquisition, project oversight.

## FIGURES

**Supplementary Figure 1.**
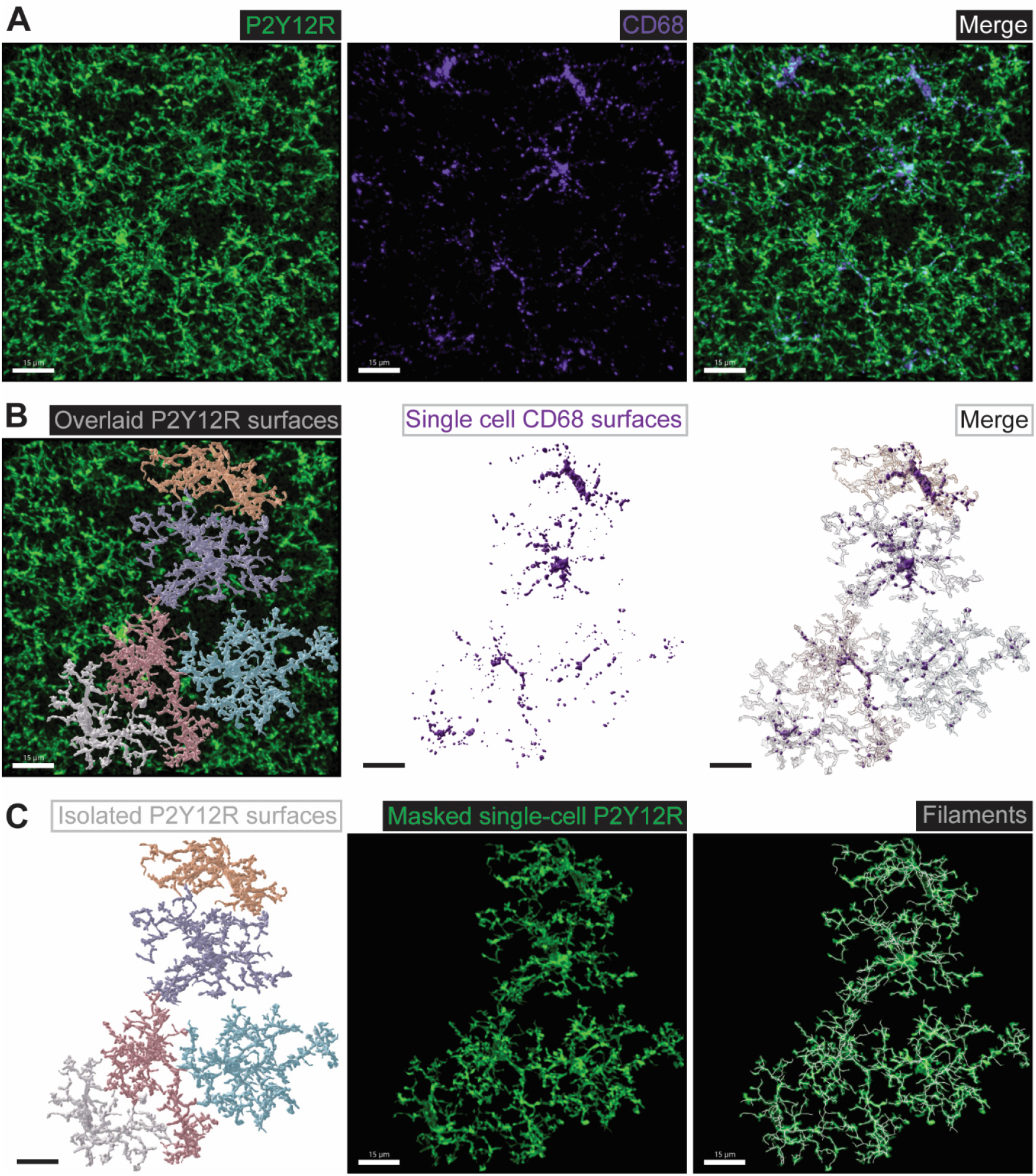
Analysis procedure for extracting morphology and phagocytic activity information from individual microglia. **A** Example full field of view from the CA1 SR of a P16 mouse. Maximum intensity projection of filtered and background subtracted (see Methods) fluorescence image of the microglia marker P2Y12R (*left*), the lysosome marker CD68 (*middle*) and a merge of the two channels (*right*). **B** Workflow for microglia phagocytic activity analysis. Extracted surfaces of individual microglia, delineated using the P12Y12R channel (*left*). A CD68 surface for CD68 located within the P2Y12R surfaces was then generated (*middle*). The volume of CD68 surface within each microglia surface is then calculated on a cell-by-cell basis (*right*). **C** Workflow for microglia morphology analysis. The individual microglia P2Y12R surfaces (*left*) were used to mask the P2Y12R fluorescence channel for each cell (*middle*). Filaments of microglia processes were constructed using these isolated microglia fluorescence images to quantify morphological parameters (*right*). For all images, scale bar is 15 µm.

